# In-silico read normalization using set multi-cover optimization

**DOI:** 10.1101/133579

**Authors:** Dilip A Durai, Marcel H Schulz

## Abstract

*De* Bruijn graphs are a common assembly data structure for large sequencing datasets. But with the advances in sequencing technologies, assembling high coverage datasets has become a computational challenge. Read normalization, which removes redundancy in large datasets, is widely applied to reduce resource requirements. Current normalization algorithms, though efficient, provide no guarantee to preserve important *k*-mers that form connections between regions in the graph. Here, normalization is phrased as a *set multi-cover problem* on reads and a heuristic algorithm, ORNA, is proposed. ORNA normalizes to the minimum number of reads required to retain all *k*-mers and their relative *k*mer abundances from the original dataset. Hence, all connections and coverage information from the original graph are preserved. ORNA was tested on various RNA-seq datasets with different coverage values. It was compared to the current normalization algorithms and was found to be performing better. It is shown that combining read error correction and normalization allows more accurate and resource efficient RNA assemblies compared to the original dataset. Further, an application was proposed in which multiple datasets were combined and normalized to predict novel transcripts that would have been missed otherwise. Finally, ORNA is a general purpose normalization algorithm that is fast and significantly reduces datasets with little loss of assembly quality.

ORNA can be found under https://github.com/SchulzLab/ORNA

## 1 Introduction

With increasing throughput and decreasing prices for modern sequencers, the generation of high coverage sequencing datasets has become routine. This has spurred the development of a number of different approaches for the *de novo* assembly of genomes and transcriptomes [18, 19] to explore the genetic diversity of life. However, assembly is a resource-intensive task.

Here we investigate how data reduction affects the performance of non-uniform RNA-seq datasets for the task of *de novo* transcriptome assembly. This is an important problem as current methods, which rely on the de Bruijn graph (DBG) for assembly, consume a lot of main memory [22, 24, 10]. However, it is also an interesting question from the perspective of information theory. What part of the data is actually being used by the assembler?

A simple approach to remove sequencing errors and reduce data size is to trim read suffixes and prefixes that are of low quality. This generally leads to decreased assembly performance [16] and does not address the high redundancy of current read datasets. Other approaches allow the efficient correction of sequencing errors in RNA-seq datasets, which generally leads to an improved assembly performance [14, 25], albeit at increased runtime because the data has to be error corrected first.

A direct approach to remove redundancy is to cluster reads according to sequence similarity and remove duplicated and highly similar reads in the clusters, *e.g.*, CD-HIT [9]. Albeit recent improvements have been made for clustering assembled transcripts [26], clustering hundred millions of reads before assembly is still challenging.

Another widely used approach, in particular in combination with assembly, has been “digital normalization” (Diginorm) [3], implemented in the khmer package [5]. Diginorm uses a min-count-sketch data structure to estimate *k*-mer abundance while streaming through the read dataset. Using a user-selected abundance threshold *t*, reads are removed once their median *k*-mer coverage goes beyond *t*. An idea similar to this is Trinity’s *in silico* normalization (TIS) which is a part of the Trinity assembler package [11]. For each read in the dataset, TIS computes the median coverage of the *k*-mers in the read. If the median coverage is less than the desired coverage, the read is always kept. Otherwise, it is kept with a probabilty which is equal to the ratio of the desired coverage and the median coverage. Recently developed NeatFreq algorithm [17] clusters the read into bins based on median *k*-mer frequency. The advantage of this kind of normalization is threefold: (i) reads with high redundancy are removed leading to reduced memory and runtime requirements for the assembly, (ii) erroneous reads may be removed as part of the process, and (iii) normalization is fast and consumes only a fraction of the memory an assembler would take. This essentially lowers the computational complexity of the assembly problem as it was shown that often a large part of the data can be removed, without significantly affecting assembly performance [3, 11]. However, previous algorithms do not give any guarantees on that important parts of the data containing useful *k*-mers would be preserved. Thus, reads that contain low-abundant but important *k*-mers may be removed. This might result in losing connections in the DBG and hence fragmented assembly. This is especially problematic for sequencing datasets with non-uniform coverage, like RNA-seq and metagenomics.

Here the Optimized Read Normalization Algorithm (ORNA) is suggested based on the idea that reads are reduced to fulfill two properties (i) the DBG backbone (unweighted nodes and edges) and (ii) relative node abundances are preserved in the reduced dataset as compared to the original DBG. It is shown that read normalization can be phrased as a *set multi-cover* optimization problem on *n* reads and we suggest a *O*(*|n| log |n|*) time heuristic algorithm that is shown to work well in practice. Analysis of normalized and error corrected data reveals that better assemblies can be produced with significant savings in runtime and memory consumption. The software is freely available at https://github.com/SchulzLab/ORNA.

## 2 Methods

### 2.1 Problem Formulation

A dataset ℛ = *{r*_1_, *r*_2_, …,*r*_*n*_*}* is a set of *n* reads where each read is a sequence of DNA bases of fixed length *s*. Each read consists of a set of short words (*k*-mers) of length *k*. Most of the *de novo* assemblers start by constructing a *de Bruijn* graph (DBG). *k*-mers obtained from all the reads in ℛ are considered as vertices. Two vertices are connected by an edge if they overlap by *k-* 1 bases. Each edge is identified by a unique label *l* of length *k* + 1, such that the source vertex is a prefix of *l* and the destination vertex is a suffix of *l*. Since the labels are also generated from the reads in the dataset, each read *r* ∈ *ℛ* can be considered as a set of *m* = *s-k* labels, i.e. *r* = (*l*_1_, *l*_2_, …, *l*_*m*_), where *l*_*i*_ is a *k* + 1-mer obtained from the *i*th position in read *r*.

The objective of a normalization algorithm is to reduce the data as much as possible without having significant impact on the quality of the assembly produced. Since an assembly is produced by traversing paths in the DBG, it may be worthwhile that a new DBG build from the normalized dataset preserves all the (unweighted) nodes and edges of the original graph. Further, normalization should maintain the relative difference of abundance between *k*-mers to resolve complex graph structures. Here, we consider a solution as Set Multi-Cover (SMC) problem. Given a dataset ℛ of *n* reads, let 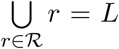 be the set of labels (*k* + 1-mers) obtained from ℛ. Every label *l ∈ L* is assigned a weight *w*_*l*_ *≥* 1. The goal of the algorithm is to find a minimum cardinality subset *ℛ′* which multi-covers *L*:

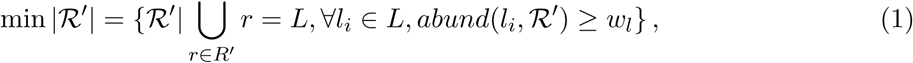

where *abund*(*l*_*i*_, *R′*) denotes the number of occurrences of *l*_*i*_ in *R′*.

The SMC problem was shown to be a NP-hard problem [4]. A common approximation approach for the SMC-problem, is the following greedy approach: an element in the universe is termed as *active* if it has not yet been covered by any of the selected sets. Cost-effectiveness of a set, to be considered for selection, is measured in terms of the number of *active* elements present in the set. The algorithm would iterate over sets and select the one which is the most cost-effective. In our application each read represents a set of labels. Thus, a data structure has to be maintained that holds the reads in an order starting from the one which has the most number of *active* labels. This order has to be updated after every iteration. For a dataset containing *n* reads, the greedy approach would take *𝒪*(*n*^2^*log n*) time assuming reads are sorted by cost-effectiveness using a binary heap or a similar data structure, which is not efficient enough for the large datasets considered here.

### 2.2 ORNA

In this work, a different heuristic algorithm based on a greedy read selection strategy is used, in which the ordering of reads based on cost-effectiveness is ignored to save runtime. The approach is summarised in algorithm 1. Set of *n* reads ℛ, a weight *w* and a *k*-mer value *k* is given as input. Since each edge label is a *k* + 1-mer, all the possible *k’* = *k* + 1-mers are obtained from the reads and are stored in a bloom filter using the *BuildBloom(ℛ,k*’) function (line 3). This functionality is implemented using the GATB library [7], which uses the *BBHash* algorithm for building a *minimal perfect hash function* [15] after counting the *k*-mers with the DSK algorithm [21]. *k*-mer counting and storing the information requires *𝒪*(*|n| log |n|*) time. A counter array *N odeCounter* is maintained for each entry in the bloom filter and is initialised to zero (line 4). This operation requires *𝒪*(*|n|*) time.

The dataset is then iterated and each read in the dataset is checked whether it contains a *k’*-mer that needs to be covered. This is done by first collecting a set *V ’* of all the *k’*-mers in the read using the *ObtainKmers*(*r, k’*) function (line 7). The counter for the *k’*-mer is incremented if its current value is less than the given weight (line 9-13). A read is accepted if it contains at-least one *k’*-mer for which the corresponding counter is incremented otherwise it is rejected (line 15-17). Steps 6-17 require *𝒪*(*|n|*) time. All the accepted reads then make the normalized dataset of ℛ_*out*_ from ℛ. The overall time complexity of the algorithm is *𝒪*(*|n| log |n|*).

An important parameter for the algorithm is the weight for each *k’* -mer. A naive way to decide the weight for each *k’*-mer is to set the same value for each. A DBG based assembler uses *k*-mer abundance information to resolve irregularities in the graph such as bubbles and tips. This information is also used for RNA-seq data to decide which transcripts are being reported. Hence, it is important to retain the relative difference in abundance between *k*-mers, which cannot be achieved with a fixed weight for all *k’*-mers. Therefore, each *k’*-mer has its own weight *w*_*k* ’_, which is defined as:

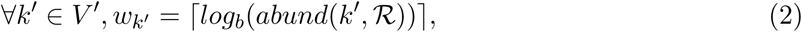

where *abund*(*k’, ℛ*) is the abundance of *k’* in ℛ. *b* is the base of the logarithm function and is given by the user.

#### algorithm 1: Set multi-cover based approach for read normalization

1. **Input:** Reads ’, weight *w*, kmer *k*
2. *k’* = *k* + 1
3. *V* = BuildBloom(*’*,*k’*)
4. NodeCounter[1…*sizeof* (*V*)] ←0
5. *R*_*out*_ = *Φ*
6. **for all** *r ∈ R* **do**
7. *V* ’ = ObtainKmers(*r*,*k’*) ⊲V‵ ⊆ *V*
8. flag=0
9. **for all** *v ∈ V ’* **do**
10. **if** *N odeCounter*(*v*) *< w* **then**
11. NodeCounter(*v*) = NodeCounter(*v*) + 1
12. flag=1
13. **end if**
14. **end for**
15. **if** *f lag* == 1 **then**
16. ℛ_*out*_ = ℛ_*out*_ *∪ r*
17. **end if**
18. **end for**
19. **Output:** Reads ℛ_*out*_

### 2.3 Data retrieval and preprocessing

Seven different datasets were used for the analyses shown in this work. Two datasets were downloaded from the SRA database-Brain dataset ([2], SRR332171) which consists of 147M paired-end (PE) reads with read length 50bps and hESC dataset ([1],SRR1020625) which has 142M PE reads with length 50bps. The datasets were error corrected, unless otherwise stated, using SEECER version 0.2 [14] with default parameters. Further, a *combined dataset* of 883M reads of length 76 bps was obtained by concatenating five ENCODE datasets: 101M PE reads from hESC (GSM758573), 192M PE reads from AG04450 (GSM765396), 207M PE reads from GM12878 (GSM758572), 165M PE reads from A549 (GSM767854) and 216M PE reads from HeLa (GSM767847).

### 2.4 Transcriptome assembly and evaluation

To analyse the quality of the assembly produced from the reduced datasets, two DBG-based *de novo* assemblers - Oases ([24], version 0.2.08) and TransABySS ([22], version 1.5.3) were used. The assemblers were run with default parameters except the *k*-mer parameter. TransABySS was run using a single *k*-mer size (*k*=21). Oases was run using multiple *k*-mer sizes (*k*=21 to *k*=49 with an increment of 2) and the assemblies obtained from all *k*-mer sizes were merged using the Oases merge-script.

The transcripts were then aligned against a reference genome using Blat ([13], version 36) and the overlap was matched against annotated Ensembl transcripts downloaded from the Ensembl database (version 65 [6]). Then the number of Ensembl transcripts that were fully assembled by at least one distinct transcript was termed as *full-length transcripts*.

For the analysis of the *combined* dataset, ORNA was used for normalization. The goal of the analysis was to determine how many transcripts are missed by assembling individual datasets, but are assembled using the *combined* datasets. All assemblies for this experiment were run with TransABySS (*k*=21). Assembly generated by using all datasets was termed *combined assembly*. Individual dataset assemblies were clustered with the *combined assembly* using CD-HIT-EST [9]. Similar sequences were clustered together (sequence identity=99%). Hence, if a transcript in the *combined assembly* is also assembled by any of the individual datasets, then it would be clustered with the sequences assembled from that dataset. All clusters which contained only the sequences from the *combined assembly* were termed as *missed clusters* and the longest sequence of the cluster was considered a *missed transcript*. Aligned missed transcripts were compared to annotations from Ensembl (version 65 [6]) and GENCODE (version 17 [12]).

### 2.5 Correlation analysis

Further, gene expression values of all RNA-seq datasets for Ensembl transcripts were obtained using Salmon (v0.8.2, [20]). The *k*-mer parameter was set to 21 for quantification. Salmon provides quantification results at transcript level. To obtain the quantification at gene level, Transcripts Per Million (TPM) values of all the transcripts for a gene were summed. Spearman correlation values between original and reduced datasets were computed using *R*.

## 3 RESULTS

### 3.1 Comparison to established normalization algorithms

The task of a read normalization algorithm is to remove the maximum number of reads without compromising the quality of the assembly produced. Most of the *de novo* assemblers start by building a DBG using a fixed/variable *k*-mer size and generate assemblies by traversing paths in the graph. In order to maintain assembly quality, ORNA was designed to: (i) retain all nodes and their connections in the DBG and (ii) retain the relative abundance difference between *k*-mers in the reduced dataset.

Diginorm and Trinity’s in-silico normalization (TIS) base their decision on the *k*-mer abundance distribution within individual reads. A read might get removed if the median abundance of *k*-mers present in the read exceed a certain threshold. Hence a *k*-mer having low abundance, if present among high abundance *k*-mers in the read, is also removed. For instance, Fig. 1 shows a toy DBG. Nodes in the graph represent *k*-mers and the abundance of each node is represented as a number inside the node. If the desired median abundance is 10 then read *r*_1_ covering node A,C,E and G would be removed since the median abundance of *k*-mers in *r*_1_ is 20. Although, this strategy may potentially remove erroneous *k*-mers, it might also result in the loss of true *k*-mers which form connections between nodes in the graph. For instance, removal of read *r*_1_ in the above example would remove the *k*-mer corresponding to node E (dashed node). Hence, the connection between node C and node G is lost, which possibly results in a fragmented assembly.

**Figure 1:**
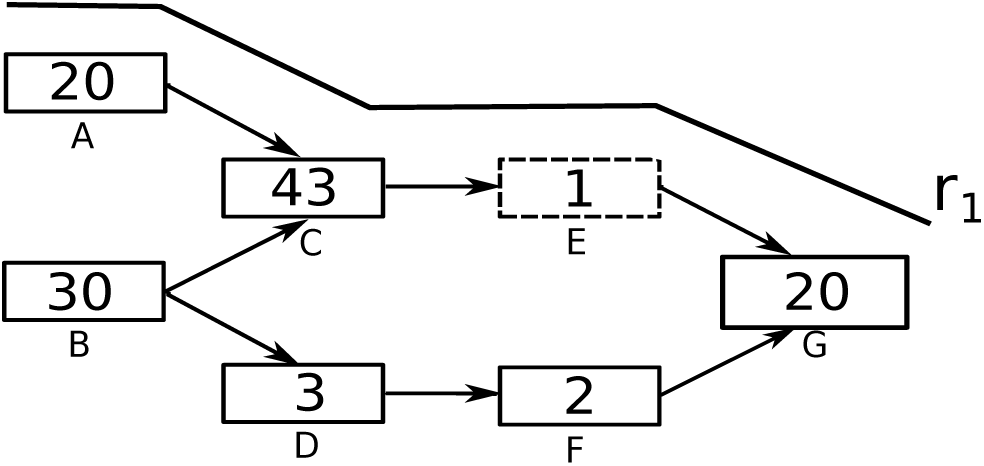
A toy DBG with one traversing read shown denoted *r*_1_. Nodes in the DBG represent *k*-mers, numbers inside nodes represent the abundance of the corresponding *k*-mer in the data. Dashed node represents the *k*-mer, which will be lost if *r*_1_ is removed.

#### ORNA retains all the kmers from the original dataset

In this work, all the assemblies used for evaluation are generated by constructing a DBG, which uses a *k*-mer size of 21. Hence the size of the edge label would be *k* + 1. Figure 2(a) shows the retention of 22-mers in reduced versions of the brain dataset by ORNA, Diginorm and TIS. Diginorm and TIS loose 2-10% of the 22-mers. ORNA considers the normalization problem as a set multi-cover problem. The set of all possible edge labels (of length *k* + 1) serves as the universe. A single read is considered as a set of *k* + 1-mers and a dataset is considered as a collection of such sets. ORNA selects the minimum number of reads, which is required to cover all the elements of the universe a certain number of times. Hence, all edge labels from the original dataset are retained.

**Figure 2:**
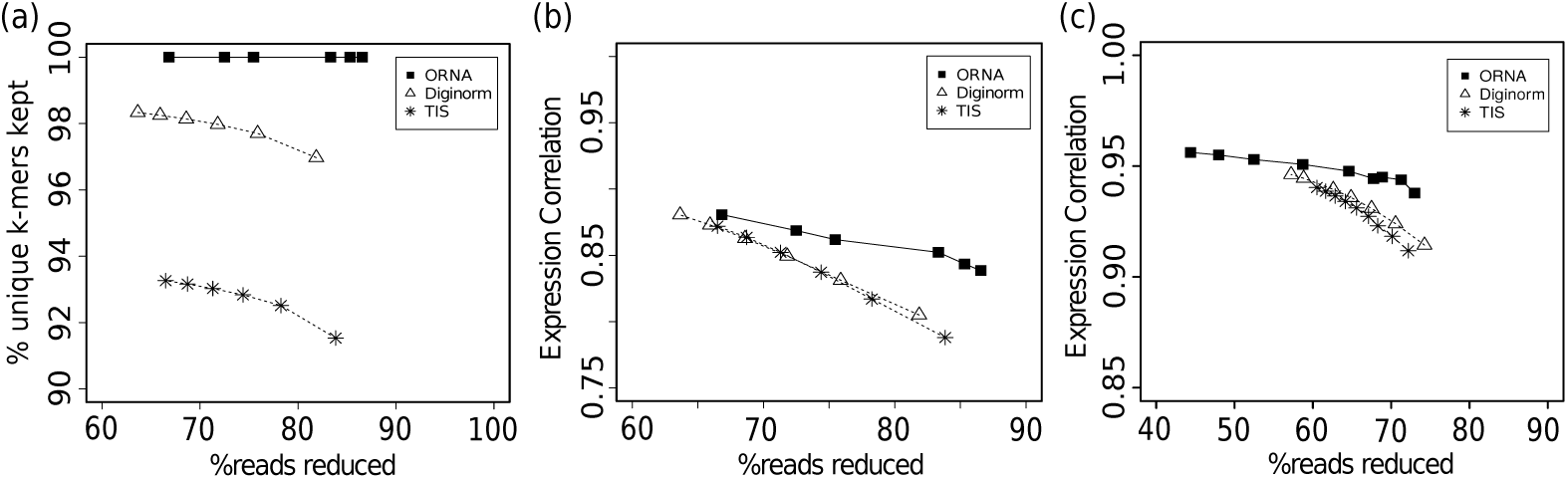
Comparison of *k*-mer information retained by ORNA, Diginorm and TIS. (a) represents percentage of unique *k*-mers retained compared to the original brain dataset (x-axis) for various levels of reduction (y-axis) by the three algorithms. (b) and (c) represent Spearman correlation values for TPM values obtained by quantifying expression of Ensembl genes using the unreduced and reduced brain dataset and hESC dataset, respectively.

#### **ORNA maintains the relative difference of abundance between** *k***-mers**

A *de novo* assembler generally uses the *k*-mer abundance information to resolve erroneous graph structures like bubbles and tips, among other things. For instance, in Fig. 1, assume that a bubble is formed by nodes C,D,E and F. An assembler would remove this by converting the less abundant path B-D-F-G to the higher abundant path B-C-E-G. Hence, it is important to maintain the relative abundance difference between the *k*-mers. ORNA uses the log_*b*_ of the abundance of the connection (*k* + 1-mer) in the original dataset. This results in large reduction of highly abundant *k*-mers and little to no reduction of lowly abundant *k*-mers, maintaining relative abundance differences. Figure 2(b) shows the comparison of spearman correlation values obtained between TPM values of the reduced and unreduced brain dataset. It can be seen that in all cases, the correlation is higher for ORNA reduced datasets as compared to using Diginorm and TIS for reduction. This indicates that for any % of reduction ORNA is able to better maintain the relative abundance of *k*-mers in genes compared to Diginorm and TIS.

#### Comparison of assembly performance

As mentioned in the above section a read normalization algorithm should not compromise on the quality of the assembly produced. But which quality measure should be used to evaluate the performance of the normalization algorithm? The number of *full-length* transcripts, as determined by aligning the assembled transcripts to a reference sequence and comparing it with the existing gene annotation, is commonly used as one of the measures and hence is also used in this work, see Methods. The total number of *full-length* transcripts obtained by running the assembler on the original unreduced dataset is considered as *complete*. The performance of a normalization algorithm is measured in terms of *% of complete*. For example, if assembling an unreduced dataset produces 2000 *full-length* transcripts and assembling a normalized dataset produces 1000 *full-length* transcripts, then it is considered that normalized data achieved *50% of complete*. A normalization algorithm *A* is better than an alternative algorithm *B*, once *A* achieves a higher *% of complete* with a similar or higher percentage of reads reduced compared to *B*.

Performance of ORNA was compared against TIS and Diginorm, with the same effective *k*-mer value=22 on two different datasets. Notably, the different parameters of the three algorithms, behave quite different with respect to the amount of reads reduced in a data-dependent manner. Therefore, the parameters of all algorithms were varied and all these results are shown in the plots, see supplement for parameter settings. Figure 3 compares the amount of read reduction (x-axis) against the assembly performance as *% of complete* (y-axis). For TransABySS (Fig. 3a,b), with *k*-mer size 21, it is observed that the quality of the assembly degrades as more reads are being removed from the dataset. All normalization algorithms perform similar at a lower percentage of reduction (60%-80% for brain and 50%-60% for hESC). But at a higher percentage of reduction (80%-90% for brain and 70%-80% for hESC), the assembly performance for Diginorm and TIS reduced datasets degrades faster than the assembly of ORNA normalized datasets. In other words, assemblies produced by ORNA reduced datasets retain equally many or more *full-length* transcripts using TransABySS.

**Figure 3:**
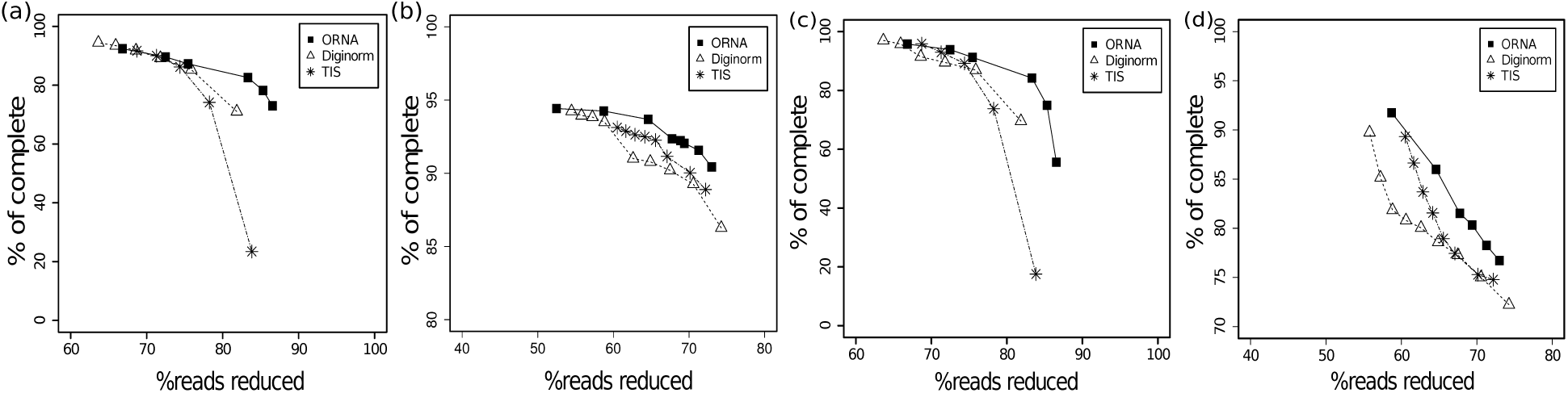
Comparison of assemblies generated from ORNA, Diginorm and TIS reduced datasets. Each point on a line corresponds to a different parametrization of the algorithms. The amount of data reduction (x-axis) is compared against the assembly performance measured as *% of complete* (y-axis, see text). (a) and (b) represent TransABySS assemblies (*k*=21) applied on normalized brain and hESC data, respectively. (c) and (d) represent Oases multi-*k*mer assemblies applied on normalized brain and hESC data, respectively.

A more challenging test is to use a multi-*k*mer based assembly strategy, where several DBGs are built for different values of the *k*-mer size. Here Oases was used to produce merged assemblies with DBGs build with *k*-mers 21-49, see Fig. 3c,d. For both the datasets a similar trend is observed as before, at higher percent of reduction (60% to 80%), ORNA assembles more *full-length* transcripts compared to Diginorm and TIS.

#### Comparison of resource requirements

ORNA stores *k*-mers in bloom filters [23] (implemented in the GATB library) making it runtime and memory efficient. Table 1 shows the comparison of memory and runtime required by ORNA against those required by Diginorm and TIS for brain, hESC and the combined dataset (see Methods) with *k*-mer value 22. For the calculations, normalized datasets were chosen to have similar read counts. ORNA and TIS were also run using one and 10 threads each. Diginorm is not parallelized. All the algorithms were run on a machine with 16GB register memory(RDIMM) and 1.5TB RAM. It can be observed that ORNA generally consumes less than half the memory and runtime required by TIS for a similar percentage of reduction. This is true for both the single and multithreaded versions. A nice property of Diginorm, is that the user can set the memory used by tuning the number of hashes and the size of each hash. A low number of hashes would reduce the runtime of Diginorm but increases the probability of false positive *k*-mers. A higher number of hashes reduces false positives but Diginorm takes longer. In this work, two different parametrizations, denoted as Diginorm^*a*^ and ^*b*^, were considered. The first uses less and the second a similar amount of memory compared to ORNA. In both cases, ORNA has an advantage of runtime over Diginorm. While Diginorm’s memory can be flexibly set, we note that ORNA uses much less space than the assembly of the reduced dataset itself, therefore not restraining the workflow.

**Table 1:** Runtime (in minutes) and memory (in GB) required by different normalization algorithms. Time and memory as obtained by running with 10 threads are shown in brackets (if possible). Note that the memory of Diginorm can be set by the user. For comparison it is set such that it uses less or similar memory than ORNA denoted as Diginorm ^*a*^ and ^*b*^, respectively. The percent of reads reduced by each method (% reduced) is shown in the first column for each dataset. The total number of reads (in millions) and the file size (in GB) of the original dataset is shown in brackets next to the dataset.

| method | brain (147M - 20.1GB) |  |  | hESC (142M - 13.2GB) |  |  | combined (883M - 98.1GB) |  |  |
| --- | --- | --- | --- | --- | --- | --- | --- | --- | --- |
|  | %reduced | time [min] | mem [GB] | %reduced | time[min] | mem [GB] | %reduced | time[min] | mem [GB] |
| ORNA | 83.3 | 104 (41) | 6.62 (5.57) | 63.90 | 58 (21) | 6.16 (6.11) | 81.4 | 740 (314) | 32.9 (33.01) |
| Diginorm<sup>a</sup> | 81.86 | 110 | 3.13 | 61.63 | 115 | 3.13 | 80.37 | 760 | 12.51 |
| Diginorm<sup>b</sup> | 81.51 | 135 | 6.26 | 62.59 | 126 | 6.26 | 79.31 | 2158 | 34.3 |
| TIS | 83.84 | 160 (95) | 20.39 (20.19) | 61.63 | 145 (127) | 13.54 (13.53) | 81.96 | 1859 (783) | 95.19 (96.09) |

### 3.2 Combined error correction and normalization generates fast and accurate *de novo* assemblies

As ORNA retains all the *k*-mers from the original dataset, useless or erroneous *k*-mers are also retained. As Fig. 3a shows, running ORNA on the corrected brain data reduces more reads compared to the uncorrected data and leads to an improvement in assembly performance, tested with Oases multi*k* assemblies. For a similar % of reduction, the error corrected data leads to more full length transcripts being assembled. This suggests that it is important to error correct the data before normalization with ORNA.

For a given dataset, assembly can be performed on - (i) the original dataset, (ii) error corrected (EC) dataset, (iii) error corrected and normalized dataset. Figures 4b and c show the maximum memory and runtimes required by applying these different strategies to the brain dataset. As seen from Figure 4b, memory required to assemble the uncorrected brain dataset is quite high. This is because of erroneous *k*-mers that complicate the graph as well as the high redundancy in the dataset. Memory is reduced by nearly 20% after error correction. It is further reduced by nearly 80% when the error corrected data is normalized using ORNA and then assembled.

**Figure 4:**
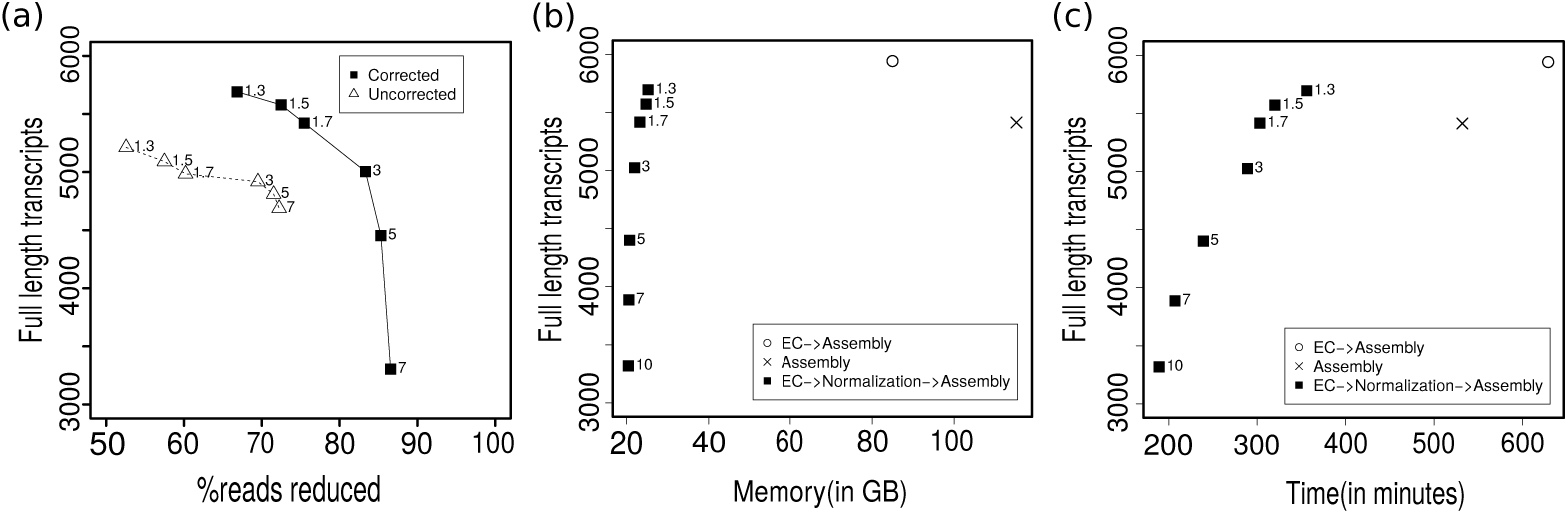
Oases assembly performance using ORNA on the brain dataset. (a) ORNA normalization on uncorrected (dashed line) and error corrected (solid line) data. The points on the curve represent how many full length transcripts (y-axis) have been assembled at a particular percent of reduction (x-axis). (b) Analysis of memory required (x-axis) by different assembly strategies namely - assembling uncorrected data (cross), assembling error corrected (EC) data (circle) and assembling error corrected and normalized data (rectangle). (c) Analysis of runtime (x-axis) required by the three strategies. The numbers next to the rectangles denote the log base *b* used for ORNA.

Runtime required to assemble a high coverage dataset is generally high (Fig. 4c). Error correcting the data improves the assembly but increases the runtime of the entire process. Normalizing error corrected data and then assembling it reduces the runtime by nearly 40% but still generates a high number of full length transcripts. Interestingly, for three ORNA parameters (*b*=1.3,1.5,1.7) the assembly quality (in terms of number of full length transcripts) is better than assembly of the uncorrected data, albeit at much lower time and memory consumption. For more stringent parameters (*b >*1.7) the quality degrades due to a high % of reduction. Note that it is likely that similar conclusions could be made using Diginorm or TIS as normalization method.

Thus in combination, error correction and normalization of RNA-seq data can be considered as potent pre-processing steps to an assembly procedure.

### 3.3 Finding novel transcripts by joint normalization of large datasets

Another important application of RNA-seq assembly methods is to find novel, unknown transcripts in well-annotated genomes by analyzing dozens of datasets of specialized tissues, *e.g.*, in cancer [27]. The common routine in these works is to assemble each RNA-seq dataset individually. It can be argued, that instead of running the assemblies on the individual datasets, a combined dataset may allow to assemble transcripts that are not well covered in an individual dataset. But combining datasets increases the size of the dataset and thereby increases the resource requirements. Hence, the dataset has to be normalized first to remove redundancy. In this work, five diverse RNA-seq datasets were concatenated to form a combined data set of 883M reads. ORNA reduced the combined dataset to 163M reads. The reduced dataset was then assembled using TransABySS. Transcripts which could only have been generated by combining the datasets were obtained and termed as *missed transcripts*, see Methods. The *missed transcripts* were aligned against the genome and compared against Ensembl annotations to estimate the number of *full-length* transcripts that have been missed by the individual assemblies. The biotypes of the *full-length* transcripts were obtained from Ensembl and GENCODE. Overall 311 missing protein coding transcripts were obtained by assembling the *combined* datasets. Along with these, 14 long non-coding RNAs, 90 non-coding RNAs and 18 pseudogenes are also part of the *missed* transcripts. Similar results may be obtained with Diginorm or TIS, albeit only ORNA guarantees that no *k*-mer information is lost.

## 4 Discussion and Conclusion

This work presents ORNA, a set multi-cover optimization-based algorithm to reduce the redundancy in NGS data without losing any *k*-mer information important for DBGs. This is done by approximating the minimal number of reads required to retain all *k*-mers from the original dataset. By generating *k*mer specific normalization weights, ORNA is able to retain the relative abundance difference between *k*-mers. This is important as various assemblers for non-uniform sequencing datasets use the abundance information to resolve complex graph substructures. ORNA, when tested on multiple datasets, is able to reduce up to 85% of the data and still able to maintain nearly 80% of useful transcripts.

The performance of ORNA was compared against two well established normalization algorithms-Diginorm and Trinity’s in-silico normalization (TIS). Brain and hESC datasets reduced by these three algorithms were assembled using Oases and TransABySS. ORNA reduced datasets were able to assemble more or equally many full length transcripts as compared to Diginorm and TIS reduced datasets. Throwing away data may show unexpected results depending on the dataset and assembler used, the same parameter of a normalization algorithm, may lead to variable reduction values. Currently, there is no clear way how to set these parameters given the data at hand. A recent work has shown how to optimize *k*mer values for multi-*k*mer assembly [8]. Such an automatic selection procedure for the *k*mer value used for normalization would be worthwhile as well.

Despite the previously shown advantage of assembly runtime and memory after read normalization, it was investigated how combining and normalizing many datasets can generate novel transcripts. Intuitively, low-coverage transcripts will never be assembled or even detected in a single sequencing run. However, once many of these experiments are combined the coverage may be sufficient for assembly. It was illustrated that combining 5 diverse datasets led to the full-length assembly of *>*400 transcripts that would have been missed otherwise. This suggests that novelty detection approaches, *e.g.* in cancer samples could use that strategy to combine and normalize all datasets.

Still, ORNA is a heuristic algorithm and can be tuned further for better results. At the moment, ORNA assumes that the data used as input is error corrected using an efficient error correcting software. As it focuses on retaining all the *k*-mers from the original dataset, any erroneous *k*-mer which escapes the filter of error correction is retained. An extra filtering criteria like the quality score can be incorporated to remove erroneous *k*-mers or to improve the ordering of the reads. Also, ORNA currently lacks support for PE reads, but extending the algorithm to PE reads is straight forward and will be added in the future.

As a conclusion, this work suggests a set multi-cover based approach for removing read redundancy in large datasets and shows an improvement over the current normalization methods, in particular for single *k*-mer transcriptome assemblies. Although ORNA was only applied on RNA-seq data, the algorithm can be also be applied for other non-uniform datasets from metagenomics or single-cell sequencing. The software is freely available at https://github.com/SchulzLab/ORNA.

## Supplemental Materials: In-silico read normalization using set multi-cover optimization

### 1 Parameters used for the evaluation

Two well established normalization methods-Diginorm and Trinity’s in-silico normalization (TIS) were used for comparing and evaluating the performance of ORNA. Given a coverage threshold *c*, Diginorm streams through the read dataset. For each read, it calculates the median abundance of *k*-mers present in the read. The abundance information is obtained from the previously accepted reads. Once the median abundance goes beyond *c*, the read is rejected.

Trinity’s in-silico normalization, on the other hand, pre-calculates the *k*-mer abundance in each read. It then iterates over the dataset. For a desired threshold *c*, TIS calculates the median abundance of the *k*-mers in the read. If the median coverage is less than the desired coverage, the read is always kept. If the median coverage is greater than the desired coverage, a random number is generated between 0 and 1. If this number is less than the ratio of the desired coverage and the median coverage, the read is kept otherwise it is removed.

**Table 1:** Parameters used different normalization algorithms. ORNA requires base *b* of the logarithm function. Diginorm and TIS requires a coverage cut-off value *c*

| Dataset | ORNA($b$) | Diginorm($c$) | TIS($c$) |
| --- | --- | --- | --- |
| SRR332171 | (1.3,1.5,1.7,3,5,7,10) | (5,10,15,20,25,30) | (5,10,15,20,25,30) |
| hESC | (1.3,1.5,1.7,3,5,7,10) | (15,20,25,30,35,45,50,55,60,65,70) | (30,35,45,50,55,60,65,70) |
| *combined* | 5 | 10 | 10 |

